# A Powerful Procedure for Pathway-based Meta-Analysis Using Summary Statistics Identifies 43 Pathways Associated with Type II Diabetes in European Populations

**DOI:** 10.1101/041244

**Authors:** Han Zhang, William Wheeler, Paula L Hyland, Yifan Yang, Jianxin Shi, Nilanjan Chatterjee, Kai Yu

## Abstract

Meta-analysis of multiple genome-wide association studies (GWAS) has become an effective approach for detecting single nucleotide polymorphism (SNP) associations with complex traits. However, it is difficult to integrate the readily accessible SNP-level summary statistics from a meta-analysis into more powerful multi-marker testing procedures, which generally require individual-level genetic data. We developed a general procedure called Summary based Adaptive Rank Truncated Product (sARTP) for conducting gene and pathway meta-analysis that uses only SNP-level summary statistics in combination with genotype correlation estimated from a panel of individual-level genetic data. We demonstrated the validity and power advantage of sARTP through empirical and simulated data. We conducted a comprehensive pathway-based meta-analysis with sARTP on type 2 diabetes (T2D) by integrating SNP-level summary statistics from two large studies consisting of 19,809 T2D cases and 111,181 controls with European ancestry. Among 4,713 candidate pathways from which genes in neighborhoods of 170 GWAS established T2D loci were excluded, we detected 43 T2D globally significant pathways (with Bonferroni corrected p-values < 0.05), which included the insulin signaling pathway and T2D pathway defined by KEGG, as well as the pathways defined according to specific gene expression patterns on pancreatic adenocarcinoma, hepatocellular carcinoma, and bladder carcinoma. Using summary data from 8 eastern Asian T2D GWAS with 6,952 cases and 11,865 controls, we showed 7 out of the 43 pathways identified in European populations remained to be significant in eastern Asians at the false discovery rate of 0.1. We created an R package and a web-based tool for sARTP with the capability to analyze pathways with thousands of genes and tens of thousands of SNPs.

**Author Summary:** As GWAS continue to grow in sample size, it is evident that these studies need to be utilized more effectively for detecting individual susceptibility variants, and more importantly to provide insight into global genetic architecture of complex traits. Towards this goal, identifying association with respect to a collection of variants in biological pathways can be particularly insightful for understanding how networks of genes might be affecting pathophysiology of diseases. Here we present a new pathway analysis procedure that can be conducted using summary-level association statistics, which have become the main vehicle for performing meta-analysis of individual genetic variants across studies in large consortia. Through simulation studies we showed the proposed method was more powerful than the existing state-of-art method. We carried out a comprehensive pathway analysis of 4,713 candidate pathways on their association with T2D using two large studies with European ancestry and identified 43 T2D-associated pathways. Further examinations of those 43 pathways in 8 Asian studies showed that some pathways were trans-ethnically associated with T2D. This analysis clearly highlights novel T2D-associated pathways beyond what has been known from single-variant association analysis reported from largest GWAS to date.

## Introduction

Genome-wide association study (GWAS) has become a very effective way to identify common genetic variants underlying various complex traits [1]. The most commonly used approach to analyze GWAS data is the single-locus test, which evaluates one single nucleotide polymorphism (SNP) at a time. Despite the enormous success of the single-locus analysis in GWAS, proportions of genetic heritability explained by already identified variants for most complex traits still remain small [2]. It is increasingly recognized that the multi-locus test, such as gene-based analysis and pathway (or gene-set) analysis, can be potentially more powerful than the single-locus analysis, and shed new light on the genetic architecture of complex traits [3, 4].

The pathway analysis jointly tests the association between an outcome and SNPs within a set of genes compiled in a pathway according to existing biological knowledge [4]. Although the marginal effect of a single SNP might be too weak to be detectable by the single-locus test, accumulated association evidence from all signal-bearing SNPs within a pathway could be strong enough to be picked up by the pathway analysis if this pathway is enriched with outcome-associated SNPs. Various pathway analysis procedures have been proposed in the literature, with the assumption that researchers could have full access to individual-level genotype data [5–9]. In practice, pathway analysis usually utilizes data from a single resource with limited sample size, as it can be challenging to obtain and manage individual-level GWAS data from multiple resources. As a result, pathway analysis often fails to identify new findings beyond what have already been discovered by the single-locus tests. To maximize the chance of discovering novel outcome-associated variants by increasing sample size, a number of consortia have been formed to conduct single-locus meta-analysis on data across multiple GWAS [10–14]. The single-locus meta-analysis aggregates easily accessible SNP-level summary statistics from multiple studies. Similarly, the pathway-based meta-analysis [15–21] that integrates the same type of summary data across participating studies could provide us a greater opportunity for detecting novel pathway associations. Future association studies focusing on identified pathways would have a much-reduced multiple-comparison burden in searching for novel variants with main or complicated nonlinear joint effects on the outcome of interest.

In this paper, we developed a pathway-based meta-analysis procedure by extending the adaptive rank truncated product (ARTP) pathway analysis procedure [9], which was originally developed for analyzing individual-level genotype data. The new procedure, called Summary based ARTP (sARTP), accepts input from SNP-level summary statistics, with their correlations estimated from a panel of reference samples with individual-level genotype data, such as the ones from the 1000 Genomes Project [22, 23]. This idea was initially used in conducting gene-based meta-analysis [24, 25] or conditional test [26]. As will be shown in the Results Section, sARTP usually has a power advantage over its competitors. In addition, sARTP is specifically designed for conducting pathway-based meta-analysis using SNP-level summary statistics from multiple studies. In real applications (e.g., the type 2 diabetes example described below), it is very common that different studies could have genotypes measured or imputed on different sets of SNPs. As a result, the sample size used in the pathway-based meta-analysis on each SNP can be quite different. Ignoring the difference in sample sizes across SNPs in a pathway-based meta-analysis would generate biased testing results.

Pathway analysis generally targets two types of null hypotheses [4], including the competitive null hypothesis [15, 16, 18–20], i.e., the genes in a pathway of interest are no more associated with the outcome than any other genes outside this pathway, and the self-contained null hypothesis [17, 21], i.e., none of the genes in a pathway of interest is associated with the outcome. The sARTP procedure focuses on the self-contained null hypothesis, as our main goal is to identify outcome-associated genes or loci. Also, as pointed out by [27], tests for the competitive null hypothesis often assume that genotype measured at different genes are independent when evaluating the association significance level. This assumption, which is generally invalid in practice, is unnecessary for sARTP when testing the self-contained null hypothesis. One may refer to [27] and [4] for more discussion and comparison of these two types of hypotheses.

The pathways defined in many public databases can consist of thousands of genes and tens of thousands of SNPs. To make the procedure applicable to large pathways, or pathways with high statistical significance, we implement sARTP with efficient and parallelizable algorithms, and adopt the direct simulation approach (DSA) [28] to evaluate the significance of the pathway association.

We demonstrated the validity and power advantage of sARTP through simulated and empirical data. We applied sARTP to conduct a pathway-based meta-analysis on the association between type 2 diabetes (T2D) and 4,713 candidate pathways defined in the Molecular Signatures Database (MSigDB) v5.0. The analysis used SNP-level summary statistics from two sources with European ancestry. One is generated from the Diabetes Genetics Replication and Meta-analysis (DIAGRAM) consortium [13], which consists of 12,171 T2D cases and 56,862 controls across 12 GWAS. The other one is based on a T2D GWAS with 7,638 T2D cases and 54,319 controls that were extracted from the Genetic Epidemiology Research on Aging (GERA) study [29, 30]. The novel T2D-associated pathways detected in the European population were further examined in Asians using summary data generated by the Asian Genetic Epidemiology Network (AGEN) consortium meta-analysis, which combined 8 GWAS of T2D with a total of 6,952 and 11,865 controls from eastern Asian populations [10].

## Material and Methods

### The Pathway-based Meta-Analysis Procedure

Here we describe the proposed method sARTP for assessing the association between a dichotomous outcome and a pre-defined pathway consisting of *J* genes. The same procedure can be applied to study a quantitative outcome with minor modifications.

#### Score Statistics and Their Variance-Covariance Matrix

We assume we have data from *L* GWA studies, with each consisting of *n*^(*l*)^ subjects, *l* = 1,⋯,*L*. Each gene in that pathway can contain one or multiple SNP(s), while any two genes may have some overlapped SNPs. For simplicity, we use superscript *l* to represent an individual study. For subject *i* in study *l*, *i* = 1,⋯,*n*^(*l*)^, let 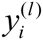 be the dichotomous outcome (e.g., disease condition, case/control status) taking values from {0,1}, and let 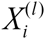 be the vector of covariates to be adjusted for. The centralized genotypes of *q* SNPs within a pathway are presented as a vector 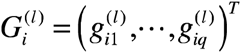 for subject *i*. We assume the following logistic regression model as the risk model

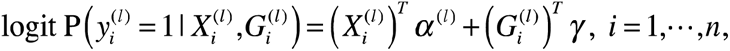

Under the self-contained null hypothesis *H*_0_ : *γ* = 0, we denote the maximum likelihood estimate of *α*^(*l*)^ as *α*̃^*l*^). Let 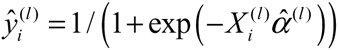 and 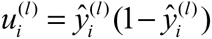.The rao’s score statistic vector on *γ*, which is the sum of score vectors from *L* participating studies, follows the asymptotic multivariate normal distribution *N*(0, *V*), where

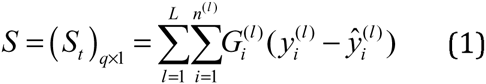

and

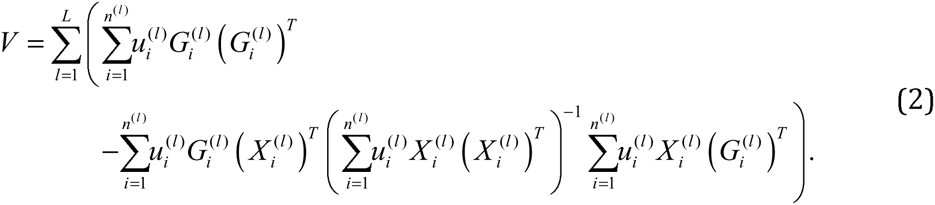

For study *l*, let 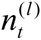 be the number of subjects having their genotypes measured as 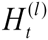 (or imputed) at SNP *t*, where 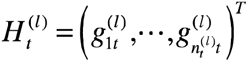. As pointed out by Hu, Berndt (24) if the covariates and genotypes are uncorrelated or weakly correlated, the covariance between scores at SNPs *t* and *S* can be approximated as

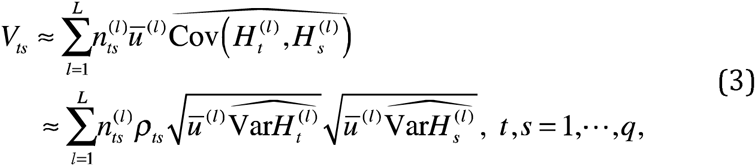

where 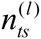 is the number of samples that have their genotypes available at both SNPs in study *l*,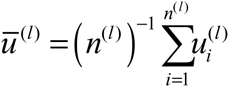 and 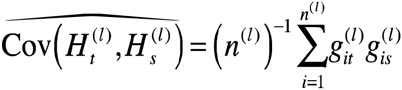.Here, we assume that the Pearson’s correlation coefficient *ρ*_*ts*_ between two SNPs is the same among all participating studies. This assumption is valid as long as subjects from all studies are sampled from the same source population, or the population under study is relatively homogeneous, such as a study of subjects with European ancestry in the United States.

When only the summary statistics, i.e., the estimated marginal log odds ratios 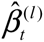 and their standard errors 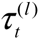 are available for each of the *L* studies, the score statistic at SNP *t*, defined by (1) can be approximated as

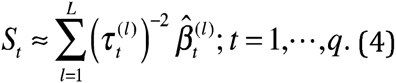

Note that 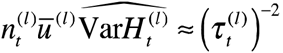,thus according to (3), we have

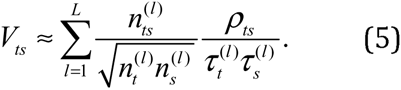

Assume that *ρ*_*ts*_ can be estimated from a public dataset (e.g., 1000 Genomes Project) and the sample sizes 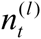 and 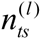 are known, we can approximately recover the variance-covariance matrix *V* = (*V*_*ts*_)_*q*×*q*_ of score statistics *S* = (*S*_*t*_)_*q*×1_. In cases when we only have the SNP p-value *p* and its marginal log odds ratio *β*̃, we can compute its standard error as 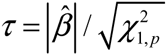 where 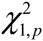 is the quantile satisfying 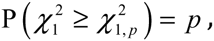, with 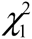 representing a 1-df chi-squared random variable.

#### Combining Score Statistics for Pathway Analysis

With recovered score statistics vector *S* and its variance-matrix *V*, we can conduct a pathway association test using the framework of the ARTP method. The ARTP method first combines p-values of individual SNPs within a gene to form a gene-based association statistic (i.e., the gene-level p-value), and then combines the gene-level p-values into a final testing statistic for the pathway-outcome association.In the original ARTP method, [9] proposed the use of a resampling-based method to evaluate the significance level of the pathway association test. Here we integrate the SNP-level score statistics into the ARTP framework and use DSA [28] to evaluate the significance level, which is much faster than the original ARTP algorithm [31]. Below is a brief summary of the improved ARTP algorithm.

First we obtain the p-values 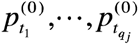 of *q*_*j*_ distinct SNPs in gene *j* as 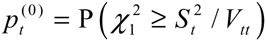. Let 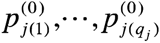 be their order statistics such that 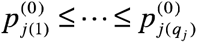. For any predefined integer *K* and SNP-level cut points *c*_1_ < ⋯ < *c*_*K*_, we define the observed negative log product statistics for that gene at cut point *c*_*k*_ as

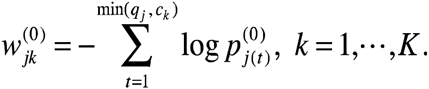

We sample *M* copies of vectors of the score statistic from the null distribution *N*(0, *V*) and convert each of them to be the tail probability of 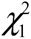 as 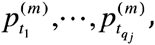, *m* = 1,⋯, *M*, which are then used to calculate 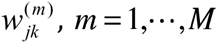. The significance of 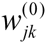 can be estimated as

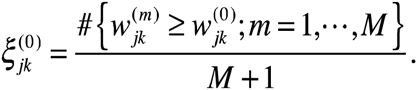

The ARTP statistic for testing association between gene *j* and the outcome is defined as 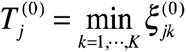. Note that for any 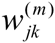, the set 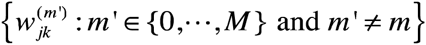 forms its empirical null distribution. The significance of 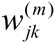 therefore can be estimated as

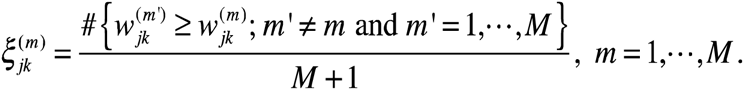

This idea, which was given by [32], can be used to avoid the computationally challenging nested two-layer resampling procedure for evaluating p-values. The p-value of 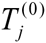 can be readily calculated as

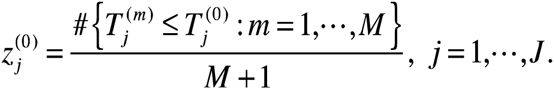

where 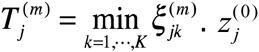 is the estimated gene-level p-value for the association between the outcome and the *j* th gene. To obtain the pathway p-value, a similar procedure as above can be applied to combine already established gene-level p-values 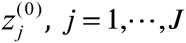, through a set of *K*' gene-level cut points *d*_1_ < ⋯ < *d*_*K′*_. For simplicity, let 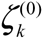 be the significance (p-value) of negative log product statistics defined on 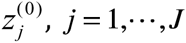, *j* = 1,⋯,*J* at specific cut point *d*_*k*_, *k* = 1,⋯,*K'*. The ARTP statistic for the pathway association is defined as 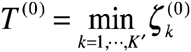. The top *d*_*k*^*^_, genes, at which 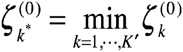, can be regarded as the set of selected candidate genes that collectively convey the strongest pathway association signal.

In the following discussion, we will use the term sARTP to represent the proposed pathway analysis procedure using the SNP-level summary statistics as input, and reserve the term ARTP to represent the original ARTP procedure that requires the individual-level genetic data. Both procedures adopt the DSA algorithm to accelerate evaluating the significance level. When performing the pathway analysis in this paper, we set SNP-level cut points as (*c*_1_,*c*_2_) = (1,2), i.e., gene-level association is summarized by one or two most significant SNPs within each gene, and gene-level cut points as *d*_*k*_ = *k* max(1,⌈*J*/20⌉), *k* = 1,⋯,10, where *J* is the number of genes in a pathway, and ⌈*J*/20⌉ is the largest integer that is less or equal to *J*/20. We used *M* = 10^5^ DSA steps to assess the significance level of each pathway in the initial screening. For pathways with estimated p-values < 10^−4^, we further refined their p-value estimetes with *M* = 10^7^ or 10^8^ DSA steps.

#### Applying sARTP to meta-analysis result

Many GWAS consortia usually publish their meta-analysis results by providing only the combined results from the fixed effects model, rather than the summary statistics from each participating study. We can apply sARTP to this meta-analysis result directly, with some modifications. First, since the reported marginal log odds ratios for each SNP by using the fixed effects inverse-variance weighting method is given by

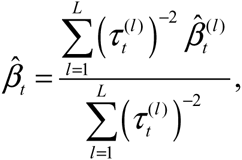

with its standard error given by

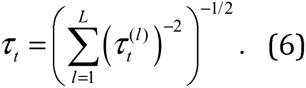

Based on (4), we can see 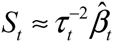. By assuming large sample sizes and certain conditions (see Appendix A), we can also approximate the covariance between *S*_*t*_ and *S*_*s*_, which is given by (5), as

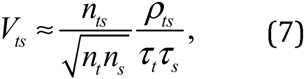

where 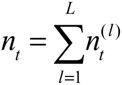, and 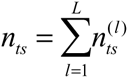. Thus, using just the meta-analysis result, without knowing summary statistics from each participating study, we can still obtain *S*_*t*_ exactly, and approximately recover *V*_*ts*_. As a result, we can carry out the pathway-based meta-analysis based on the SNP-level meta-analysis result as if it were summary data from a single study. We call this approach the Meta-analysis based sARTP (MsARTP).

However, to apply the MsARTP, we need additional sample size information *n*_*t*_ and *n*_*st*_ in order to properly estimate the variance-covariance matrix defined by (7). If the same set of SNPs are studied by all participating studies, we have *n*_*t*_ = *n*_*s*_ = *n*_*st*_, and the approximation (7) becomes 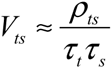, i.e., we can obtain the estimated variance-covariance matrix without knowing *n*_*t*_ and *n*_*st*_. But in most applications, not all GWAS choose the same SNP genotyping array, even after the imputation using the same reference genomes. As a result, the SNP coverage, i.e., the set of SNPs evaluated in each participating study can be quite different. In those situations, we need to know the SNP coverage information in each participating study in order to obtain *n*_*s*_ and *n*_*st*_. We will show in the Results Section that using MsARTP with an inappropriate uniform coverage assumption (i.e., *n*_*ts*_ = *n*_*t*_ = *n*_*s*_), which is commonly made by many multi-locus approaches, can lead to inflated type I error.

Given SNP-level summary statistics from each participating study, we can either apply sARTP directly, or first conduct a SNP-level meta-analysis, and then apply MsARTP to the meta-analysis result. These two approaches use the same score statistics, and different but consistent estimates for the variance-covariance matrix. Numeric experiments in the Results Section suggest that these two approaches generate vary similar pathway p-values.

### Study Materials

#### Pathway and Gene Definition

We downloaded definitions for 4,716 human and murine (mammalian) pathways (gene sets) from the MSigDB v5.0 (C2: curated gene sets). Genomic definitions for genes were downloaded from Homo sapiens genes NCBI36 and reference genome GRCh37.p13 using the Ensemble BioMart tool.

#### DIAGRAM Study

The DIAGRAM (DIAbetes Genetics Replication And Meta-analysis) consortium conducted a large-scale GWAS meta-analysis to characterize the genetic architecture of T2D [13]. We downloaded the summary statistics generated by the DIAGRAMv3 (Stage 1) GWAS meta-analysis from www.diagram-consortium.org [13]. The meta-analysis studied 12 GWAS with European ancestry consisting of 12,171 cases and 56,862 controls. Up to 2.5 million autosomal SNPs with minor allele frequencies (MAFs) larger than 1% were imputed using CEU samples from Phase II of the International HapMap Project. Study-specific covariates were adjusted in testing T2D-SNP association under an additive logistic regression model [13]. SNP-level summary statistics from each GWAS were first adjusted for residual population structure using the genomic control (GC) method [33], and then combined in the fixed effects meta-analysis.

We sorted 2.5 million autosomal SNPs by their corresponding meta-analysis sample sizes in Figure S1, which shows that there are two major groups of SNPs with equal sample sizes. One group of 469,985 SNPs (19.0%) had 12,171 cases and 56,862 controls, which included all the available samples in the meta-analysis; another group of 1,431,361 SNPs (57.9%) had 9,580 cases and 53,810 controls. Since the calculation of covariance *V*_*ts*_ in (7) relies on *n*_*ts*_, the number of samples having genotypes available at both SNP _s_ and SNP *t*, in order to obtain an accurate estimate of *n*_*ts*_, we focused on these two groups of SNPs, which in combination had a total of 1,901,346 SNPs. For any two SNPs in this reduced set, it is certain *n*_*ts*_ = min(*n*_*t*_, *n*_*s*_). The Pearson’s correlation coefficients *ρ*_*ts*_ were estimated using an external reference panel consisting of genotypes on 503 European subjects (CEU, TSI, FIN, GBR, and IBS) from the 1000 Genomes Project (Phase 3, v5, 2013/05/02).

#### GERA Study

We assembled a GWAS on T2D from the Genetic Epidemiology Research on Adult Health and Aging (GERA, dbGaP Study Accession: phs000674.v1.p1). The GERA project includes a cohort of over 100,000 adults who are members of the Kaiser Permanente Medical Care Plan, Northern California Region, and participating in the Kaiser Permanente Research Program on Genes, Environment, and Health (RPGEH). From the GERA data, we compiled a GWAS with 7,638 T2D cases and 54,319 controls (subjects without T2D) who self-reported to be non-Hispanic White Europeans in the RPGEH survey. We performed the genotype imputation with IMPUTE2 [34] using CEU reference samples from Phase II of the International HapMap Project. After removing SNPs with low imputation quality (*r*^2^ < 0.3), we ended up with 2.4 million SNPs for further analysis. In the single-locus analysis, we adjusted for the categorized body mass index (BMI) provided in the downloaded dataset (adding a category for missing BMI), gender, year of birth (in five-year categories), a binary indicator on whether or not a participant was diagnosed with cancer (includes malignant tumors, neoplasms, lymphoma and sarcoma), and the top five eigenvectors for the adjustment of population stratification. In the following discussion, we refer this assembled T2D GWAS as the GERA study.

When analyzing the SNP-level summary data from the GERA study, the Pearson’s correlation coefficients *ρ*_*ts*_ were estimated using an external reference panel consisting of genotypes on 503 European subjects from the 1000 Genomes Project.

#### AGEN-T2D Study

The Asian Genetic Epidemiology Network (AGEN) consortium carried out a meta-analysis by combining eight GWAS of T2D with a total of 6,952 cases and 11,865 controls from eastern Asian populations [10]. The meta-analysis was conducted with the fixed effect model. We obtained SNP-level summary statistics on 2.6 million imputed and genotyped autosomal SNPs from AGEN, and used this summary data to evaluate whether pathway associations identified in European populations remain to be present in Asians. We adopted an external reference panel consisting of 312 eastern Asian subjects (103 from CHB, 105 from CHS, and 104 from JPT) from the 1000 Genomes Project for the variance-covariance matrix estimation in the pathway analysis.

## Results

### Simulation Studies

Firstly, we conducted a simulation study to evaluate the empirical size of sARTP and MsARTP. Secondly, we compared empirical powers of different strategies for carrying out pathway-based meta-analysis that integrated summary statistics from multiple studies. We also evaluated whether results from sARTP were consistent with the ones from MsARTP. Thirdly, we compared our method to the recently developed method aSPUsPath [8] that can be used for pathway-based meta-analysis. We used the R package, aSPU (version 1.39), with the default settings given in [8,17] to conduct the aSPUsPath test.

#### Empirical Size of sARTP and MsARTP

To evaluate the empirical size of sARTP and MsARTP, we conducted a simulation study by using individual-level GWAS data of the pathway PUJANA_BREAST_CANCER_WITH_BRCA1_MUTATED_UP (including 728 SNPs in 50 genes) from the GERA study. We picked 12,000 samples randomly for this experiment. By keeping their genotypes unchanged, we randomly assigned 6,000 subjects as cases and the remaining as controls to generate 500,000 datasets. We split each dataset into three case-control studies, each with 2,000 cases and 2,000 controls. To mimic the scenario when not all studies have their genotypes measured on the same set of SNPs (such as the one occurred in the DIAGRAM and AGEN data), we assumed that each case-control study had genotypes measured on only half of SNPs in the pathway. For each generated dataset that consisted of three case-control studies, we applied sARTP to the SNP-level summary data obtained from each case-control study, and MsARTP to the meta-analysis result based on the three case-control studies, with the variance-covariance matrix estimated by an external reference panel (with 503 European reference samples from the 1000 Genomes Project), or an internal reference panel (with 500 samples randomly selected from the GERA data).

Based on results from the 500,000 generated datasets, this simulation study showed that both sARTP and MsARTP, using the internal or external reference samples, can well control their empirical sizes (Table 1). Given the same reference panel, the p-values estimated from sARTP and MsARTP are highly consistent (Pearson’s correlation coefficient > 0.99). Furthermore, the p-values of sARTP (or MsARTP) estimated with an external or internal reference panel are also very consistent (Pearson’s correlation coefficient > 0.99). More numeric experiments demonstrating the validity of sARTP under the null are described in Appendix B.

**Table 1:** Empirical sizes of the sARTP, MsARTP, and MsARTP-u procedures

|  |  | Size |  |  |  |  |
| --- | --- | --- | --- | --- | --- | --- |
| Reference |  | 0.05 | 0.01 | 0.005 | 0.001 | 0.0005 |
| sARTP | External <sup>a</sup> | 0.050 | 0.0093 | 0.0040 | 0.00078 | 0.00044 |
|  | Internal <sup>b</sup> | 0.046 | 0.0087 | 0.0042 | 0.00074 | 0.00040 |
| MsARTP | External | 0.048 | 0.0093 | 0.0040 | 0.00076 | 0.00041 |
|  | Internal | 0.048 | 0.0084 | 0.0042 | 0.00074 | 0.00041 |
| MsARTP-u | External | 0.082 | 0.018 | 0.0081 | 0.0013 | 0.00064 |
|  | Internal | 0.094 | 0.022 | 0.011 | 0.0016 | 0.00081 |

Empirical sizes are estimated based on 500,000 datasets simulated from the GERA data.
<sup>a</sup>Using 503 European samples from the 1000 Genomes Project as an external reference;
<sup>b</sup>Using 500 samples from the GERA data as an internal reference.

To demonstrate the importance of knowing *n*_*t*_ and *n*_*ts*_ when applying MsARTP to the meta-analysis result, we analyzed each simulated dataset using MsARTP assuming the uniform coverage (*n*_*ts*_ = *n*_*t*_ = *n*_*s*_). We called this approach MsARTP-u. It is clear from Table 1 that MsARTP-u assuming the uniform coverage suffers from inflated type I errors with either the internal or external reference panel.

#### Empirical Power of sARTP and MsARTP for Pathway-based Meta-analysis

We conducted a set of simulation studies to compare the power of different strategies to carry out pathway analysis when SNP-level summary statistics were available from multiple studies. We considered a hypothetical pathway consisting of 50 genes randomly selected from chromosome 17, each with 20 randomly chosen SNPs. The joint genotype distribution at the 20 SNPs within each gene was defined by the observed genotypes in the GERA study. We further assumed that all genes in that pathway are independent. This assumption is unnecessary for sARTP and MsARTP, but it was introduced for simplifying the simulation. For the risk model, we assumed the first *M* (*M* = 5,10,15) genes were associated with the outcome. Within each outcome-associated gene, we picked the SNP with its MAF closest to the median MAF level within the gene to be functional. We considered the following risk model

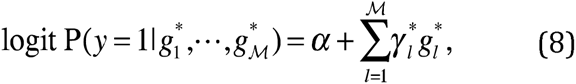

where 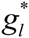 is the genotype (encoded as 0,1, or 2 according to counts of minor alleles) at the functional SNP within gene *l*. Under this model, 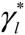 is also the marginal log odds ratio for the *l*th functional SNP [9]. Given the sample sizes of cases and controls, and the MAF of the *l*th functional SNP, 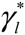 was chosen such that the theoretical power of the trend test to detect the *l*th functional SNP is equal to ***P*** (***P*** = 0.3,0.4), with 0.05 as the targeted type I error rate. For every pair of (***M**,**P***), we generated 1,000 datasets, each consisting of three case-control studies, with the same sample size and SNP coverage configurations used for evaluating the empirical size. Given the genotype distribution in the general population, individual-level genotype data for a case-control study can be generated according to the assumed risk model (8).

We assumed that only SNP-level summary statistics from each of the three studies were available. For each simulated dataset, we applied sARTP and MsARTP, using either an internal or external reference panel to estimate the variance-covariance matrix. The sARTP and MsARTP approaches integrate association evidence across SNP-level summary statistics, which are obtained by pooling information from all participating studies on individual SNPs. As a comparison, we also considered a naïve approach, in which we first applied sARTP to analyze the summary statistics from each study separately, and then combined the three pathway p-values with Fisher’s method. This naïve approach could be useful when the researchers do not have access to the SNP-level summary data but the pathway p-values from individual studies. The empirical powers are compared at the type I error level of 0.05, and are summarized in Table 2. It is obvious that the pathway-based meta-analysis using sARTP, with either the internal or external reference panel, have almost the same level of power as the MsARTP method. It is also evident that both sARTP and MsARTP are more powerful than the naïve approach, which suggests that it is always be beneficial to have the SNP-level summary statistics from each participating study, or SNP-level meta-analysis result when conducting a pathway analysis.

**Table 2:** Power comparisons under the type I error rate of 0.05 when analyzing data from three studies

| $\mathcal{P}^a$ | $\mathcal{M}^b$ | Internal reference <sup>c</sup> | | | External reference <sup>d</sup> | | |
| --- | --- | --- | --- | --- | --- | --- | --- |
|  |  | sARTP | MsARTP | Fisher | sARTP | MsARTP | Fisher |
| 0.3 | 5 | 0.165 | 0.170 | 0.110 | 0.170 | 0.167 | 0.105 |
|  | 10 | 0.405 | 0.402 | 0.229 | 0.399 | 0.401 | 0.221 |
|  | 15 | 0.573 | 0.578 | 0.334 | 0.564 | 0.561 | 0.323 |
| 0.4 | 5 | 0.292 | 0.293 | 0.162 | 0.295 | 0.297 | 0.154 |
|  | 10 | 0.642 | 0.637 | 0.363 | 0.640 | 0.635 | 0.362 |
|  | 15 | 0.858 | 0.858 | 0.574 | 0.855 | 0.856 | 0.561 |

For every pair of $\mathcal{P}$ and $\mathcal{M}$ , the empirical powers are computed from 1,000 simulated datasets at the level of 0.05. Each dataset contains three studies. The pathway consists of 50 independent genes, each with 20 SNPs. Fisher's method is used to combine the three pathway p-values obtained by applying sARTP to the SNP-level summary data from each of three studies separately.
<sup>a</sup> The theoretical power of the single-locus trend test on the functional SNP under the type I error rate of 0.05, given the sample sizes of cases and controls, and the MAF of the functional SNP;
<sup>b</sup> The number of genes including the functional SNPs;
<sup>c</sup> Using 500 samples from the GERA data as an internal reference;
<sup>d</sup> Using 503 European samples from the 1000 Genomes Project as an external reference.

Given the SNP coverage information, the MsARTP method is a valid pathway association test that has well controlled type I error and similar power to the sARTP method. In the following analysis, either sARTP or MsARTP is chosen depending on the type of available data. For the sake of simplicity, we always label the chosen procedure as sARTP.

#### Power Comparison between sARTP and aSPU sPath

Since the aSPUsPath method in the current aSPU package cannot handle summary data from multiple studies, or meta-analysis results from studies with varied SNP coverage, we focused on the scenario with just one study, and adopted the similar simulation strategy as the one used by [8] to compare the power between sARTP and aSPUsPath. We simulated haplotypes on a set of SNPs within a gene in the general population using the algorithm of Wang and Elston (35). Then the joint genotypes on a subject can be formed by randomly pairing two haplotypes. In brief, we first chose the MAF for each SNP by randomly sampling a value from the uniform distribution *U*(0.1,0.4). Then for the set of SNPs in a gene we sampled a latent vector *Z* = (*z*_1_, ⋯, *Z*_*q*_)^*T*^ from a multivariate normal distribution with a covariance matrix Cov(*z*_*i*_, *z*_*j*_) = *ρ*^|*i*-*j*|^,1≤*i*,*j*≤*q*, where *ρ* was sampled from the uniform distribution *U*(0, 0.8) for a given gene. We randomly picked 50% of the SNPs and converted their simulated *z*_*i*_ into minor and major alleles (coded as 0, 1), with the cuts chosen for each *z*_*i*_ such that the resultant minor allele has its frequency defined by the specified MAF. For the remaining SNPs, we used the same algorithm to dichotomize –*z*_*i*_ into minor and major alleles. This created a more realistic haplotype structure such that a haplotype can consist of a mixture of minor and major alleles. Genotypes on SNPs from different genes were generated independently.

Given the number of genes (20, 50, or 80) in a pathway, the proportion of genes (5%, 10%, 20%, and 30%) associated with the outcome, and a chosen common value for all log odds ratios (*γ*^*^) in the risk model (8), we repeated the following steps to generate 1,000 case-control studies, with each consisting of 1,000 cases and 1,000 controls. First, the number of SNPs within each gene was randomly chosen from 10 to 100. Second, for each randomly selected outcome-associated gene, we randomly picked a functional SNP. Third, we use the aforementioned algorithm of Wang and Elston (35) to generate the individual-level genotype data for a case-control study according to the specified risk model. We also considered the situation where all *γ*^*^ in the risk model (8) had the same magnitude but different directions. More precisely, when generating a case-control study at the third step, we defined the risk model (8) by randomly choosing the direction of each log odds ratio to be positive or negative with equal probability. Furthermore, we considered a more complex scenario where each outcome-associated gene had one or two functional SNPs, each with equal probability.

All simulation results are given in Table S1 and Table S2. It is clear that sARTP are generally more powerful than aSPUsPath, especially when the signal-to-noise ratio (the proportion of genes including a functional SNP) is relatively low. The two types of tests tend to have comparable performance when the signal-to-noise ratio increases to 30%, although it is uncommon for a candidate pathway to have such a high signal-to-noise ratio in real applications. For example, among the 4,713 candidate pathways analyzed in the next section, only 4.2% and 0.9% of the pathways have over 20% and 30% of their genes that are likely to contain association signals (i.e., with gene-level p-values < 0.05).

From Table SI and Table S2, we also notice that the advantage of sARTP over aSPUsPath is more evident if not all minor alleles of the functional SNPs are deleterious (or protective) variants (i.e., *γ*_*_ in the risk model (8) are not all positive). This is expected, as the sARTP approach does not take the effect direction of the minor allele at each SNP into consideration, while aSPUsPath integrates a set of candidate statistics, including the one similar to the burden test that assumes all minor alleles are either deleterious or protective. When this assumption is not valid, the inclusion of the burden test statistic in aSPUsPath is unlikely to enhance the power, but certainly would increase the multiple-testing penalty.

### Evaluation of sARTP using Data from T2D Studies

To demonstrate the consistency between results obtained by sARTP using SNP-level summary statistics and the ones by ARTP using individual-level genotype data, we compared pathway analysis results from three different procedures on the 4,713 candidate pathways using the GERA GWAS data. Details on how those 4,713 pathways were pre-processed are given in the Results of T2D Pathway Analysis Section. We applied sARTP to the SNP-level summary statistics generated from the GERA study, using either an internal or an external reference panel. We also obtained the pathway p-values by directly applying the ARTP method to the individual-level GERA GWAS data. Figure 1 shows the comparison among p-values from these three analyses, and demonstrates that all three approaches can generate very consistent results.

**Figure 1.**
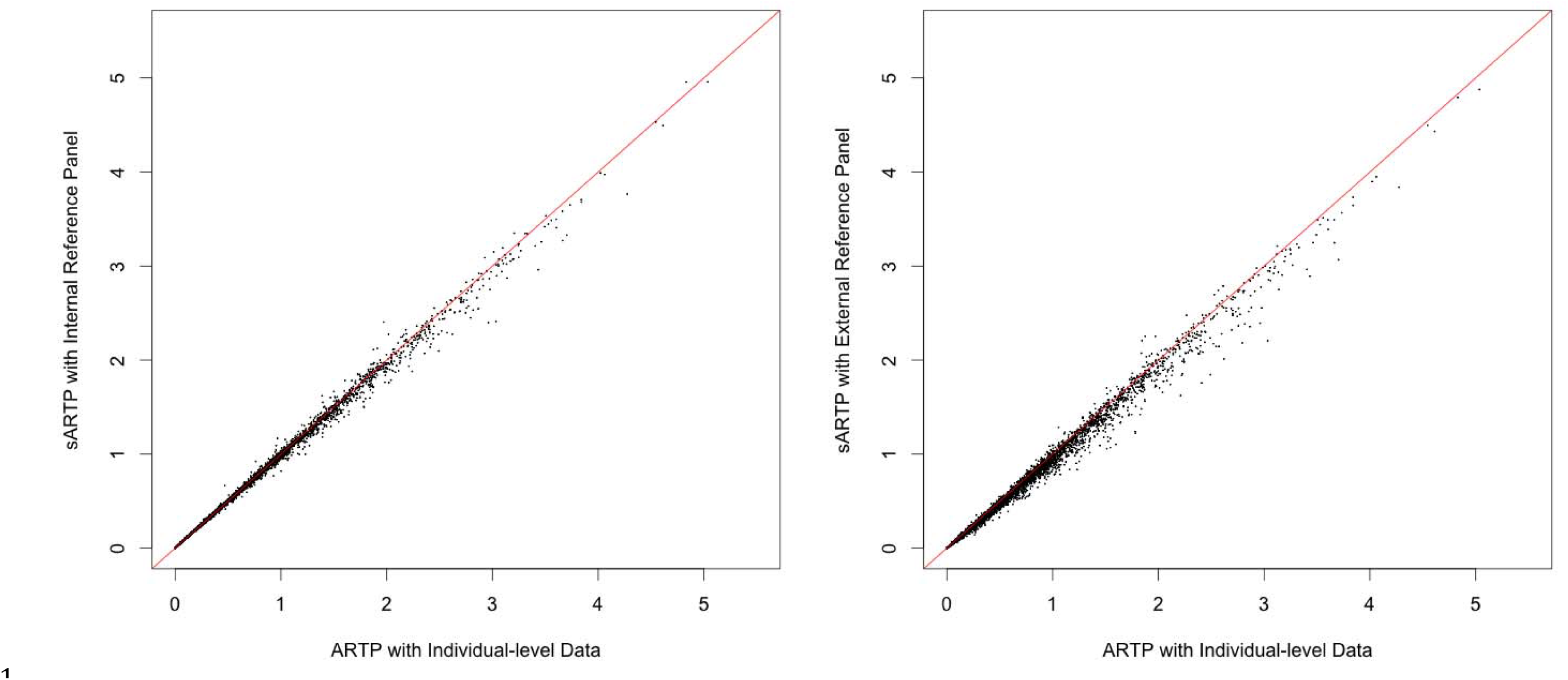
Comparisons of p-values from three types of pathway analyses on the GERA data. Based on the GERA data, 4,713 pathways are analyzed in three different ways. Pathway p-values obtained by ARTP using the GERA individual-level genetic data (x-axis) are compared with the ones obtained by sARTP using summary statistics in combination with the internal reference panel that consists of 500 randomly selected GERA samples (left), and the ones using the summary statistics in combination with the external reference panel that consists of 503 European subjects from the 1000 Genomes Project (right).

### Results of T2D Pathway Analysis

#### Findings from the European Populations

Since our goal was to identify new susceptibility loci for T2D through the pathway analysis, we excluded 170 high evidence T2D associated SNPs that were either listed in [13] or found from the GWAS Catalog satisfying the following three conditions simultaneously: (1) were investigated by GWAS of samples with European ancestry; (2) had reported p-values < 10^−7^ on the initial study; and (3) were replicated on independent studies. We excluded 195 SNPs that has their single-locus testing p-values less than 10^−7^ in either DIAGRAM or GERA data to ensure that the pathway analysis result was not driven by a single SNP. In addition, we further excluded genes within a ±500kb region from each of the removed SNPs to eliminate potential association signals that could be caused by linkage disequilibrium (LD) with the index SNPs.

We conducted three types of pathway-based meta-analyses using sARTP, including the one using the DIAGRAM SNP-level summary statistics, the one using the GERA SNP-level summary statistics, and the pathway meta-analysis combining SNP-level summary statistics from both DIAGRAM and GERA studies. When applying the pathway-based meta-analysis to a single gene, we refer to this as the gene-level meta-analysis. We used the external reference panel of 503 Europeans from the 1000 Genomes Project to estimate the variance-covariance matrix.

Before performing a pathway analysis, we applied LD filtering to remove redundant SNPs. For any two SNPs with their pairwise squared Pearson’s correlation coefficient > 0.9 estimated from the external reference panel from the 1000 Genomes Project, we removed the one with a smaller value defined as, 2*f*(1 – *f*)*n*_0_*n*_1_*n*^−1^, where *n*_0_ and *n*_1_ are numbers of controls and cases, *n* = *n*_0_ + *n*_1_, and *f* is the MAF based on the reference panel. This value is proportional to the non-centrality parameter of the trend test statistic at a given SNP. We also excluded SNPs with MAF <1%. After all SNP filtering steps, we had a total of 4,713 pathways for the analysis. The summary of the number of genes and SNPs used in each pathway analysis is given in Figure S2.

The DIAGRAM study had a genomic control inflation factor *λ*_*GC*_ = 1.10 based on the published meta-analysis result. The assembled GERA T2D GWAS had *λ*_*GC*_ = 1.08. When conducting the pathway analysis on each of two studies, we adjusted the inflation by using the corresponding 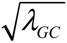 to rescale the standard error of estimated log odds ratio at each SNP. The single-locus meta-analysis combining results from DIAGRAM and GERA datasets had an inflation factor *λ*_*GC*_ = 1.067 after each study had adjusted for its own inflation factor. We further adjusted this inflation in the pathway and gene-level meta-analysis when combining SNP-level summary statistics from both studies using formulas (4) and (5).

The Q-Q plots of gene-level and pathway p-values are given in Figure 2. Gene-level p-value Q-Q plots based on the three analyses show no sign of inflation with their *λ*_*GC*_ close to 1.0, but suggest that there are enriched gene-level association signals at the tail end. The pathway p-value Q-Q plots, on the other hand, shift away from the diagonal identify line and have much higher *λ*_*GC*_, which suggests that T2D associated genes are preferably included in pathways under study. In fact, it can be seen from Figure S3 that a gene with a smaller gene-level meta-analysis p-value tends to be included in more pathways, even though the 4,713 pathways collected from MSigDB v5.0 are not specifically defined for the study of T2D.

**Figure 2.**
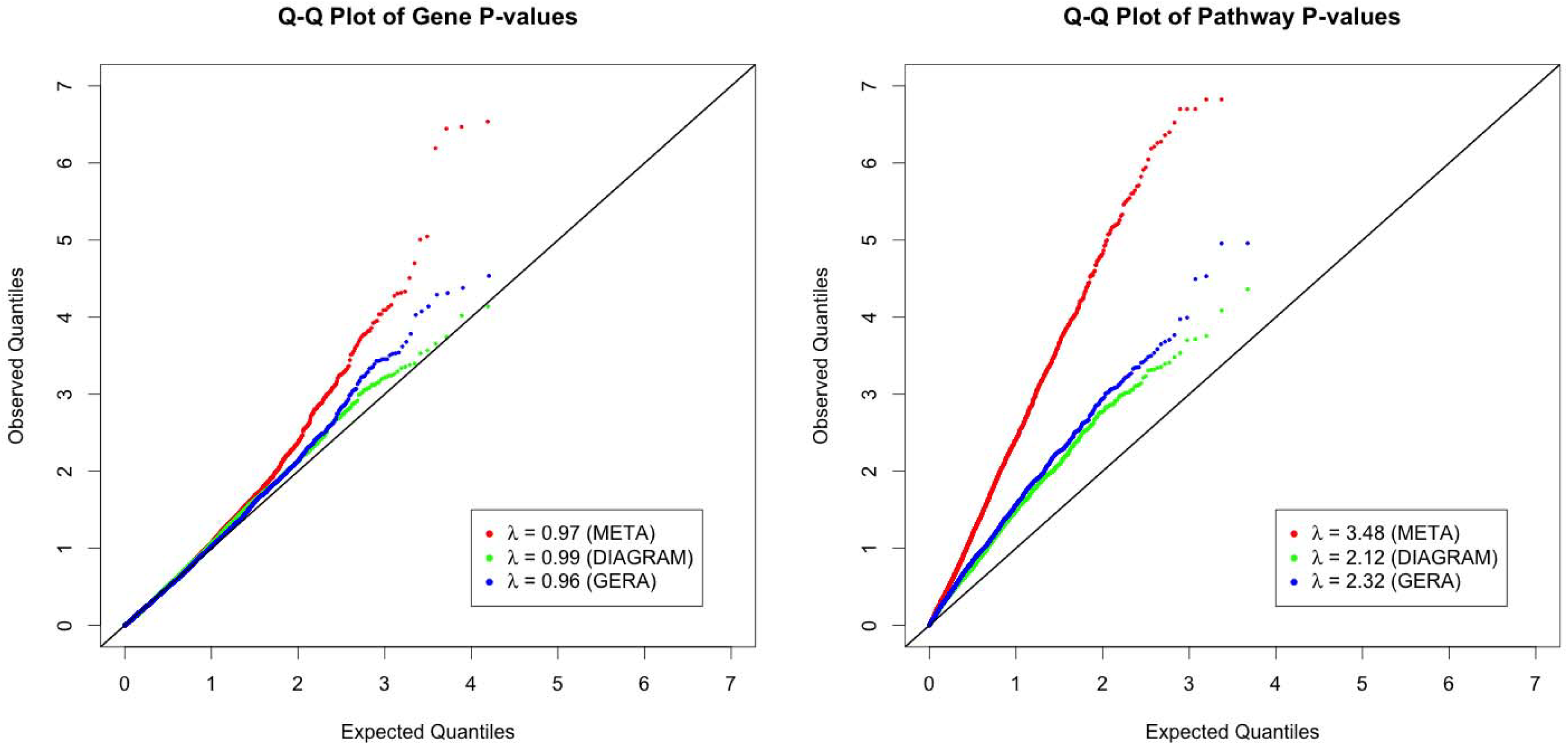
Q-Q plots of gene-level and pathway p-values based on the sARTP procedure on the DIAGRAM study, the GERA study, and the two studies combined. Left: Q-Q plots of gene-level p-values on 15,946 genes based on the sARTP gene-based analysis of the DIAGRAM study (DIAGRAM), the GERA study (GERA), and the two studies combined (META)Right: Q-Q plots of pathway p-values on 4,713 pathways based on the sARTP pathway analysis of the DIAGRAM study (DIAGRAM), the GERA study (GERA), and the two studies combined (META)

Figure 2 illustrates that the gene and pathway level signal from the GERA study tends to be slightly stronger than that from the DIAGRAM study. The main reason is that the DIAGRAM summary result had gone through two rounds of inflation adjustments, with the first round done at each participating study, and the second round on the meta-analysis result. Also, its second round adjustment (*λ*_*GC*_ = 1.10) is larger than the one applied to the GERA study (*λ*_*GC*_ = 1.08). Adjusting for *λ*_*GC*_ in the pathway analysis could be too conservative, since some proportion of the inflation can be caused by the real polygenic effect. A less conservative adjustment could be possible, but it might not be adequate. More discussions on this issue are given in the Discussion Section.

Based on the pathway meta-analysis on a total of 4,713 pathways, we identified 43 significant pathways with p-values less than 1.06×10^−5^, the family-wise significant threshold based on the Bonferroni correction. Their pathway meta-analysis results as well as results from individual studies are summarized in Table 3. More detailed results on each of 43 significant pathways are given in the Figures S6-S48, and Supplemental Data. There are a total of 15,946 unique genes in all 4,713 pathways. The top 50 genes with smallest gene-level p-values based on the gene meta-analysis are listed in Table S3. Because of the LD filtering, a gene belonging to two pathways might end up with slightly different sets of SNPs. To remove this ambiguity, we obtained the gene-level p-values by conducting a gene-level meta-analysis on each gene separately.

**Table 3.**
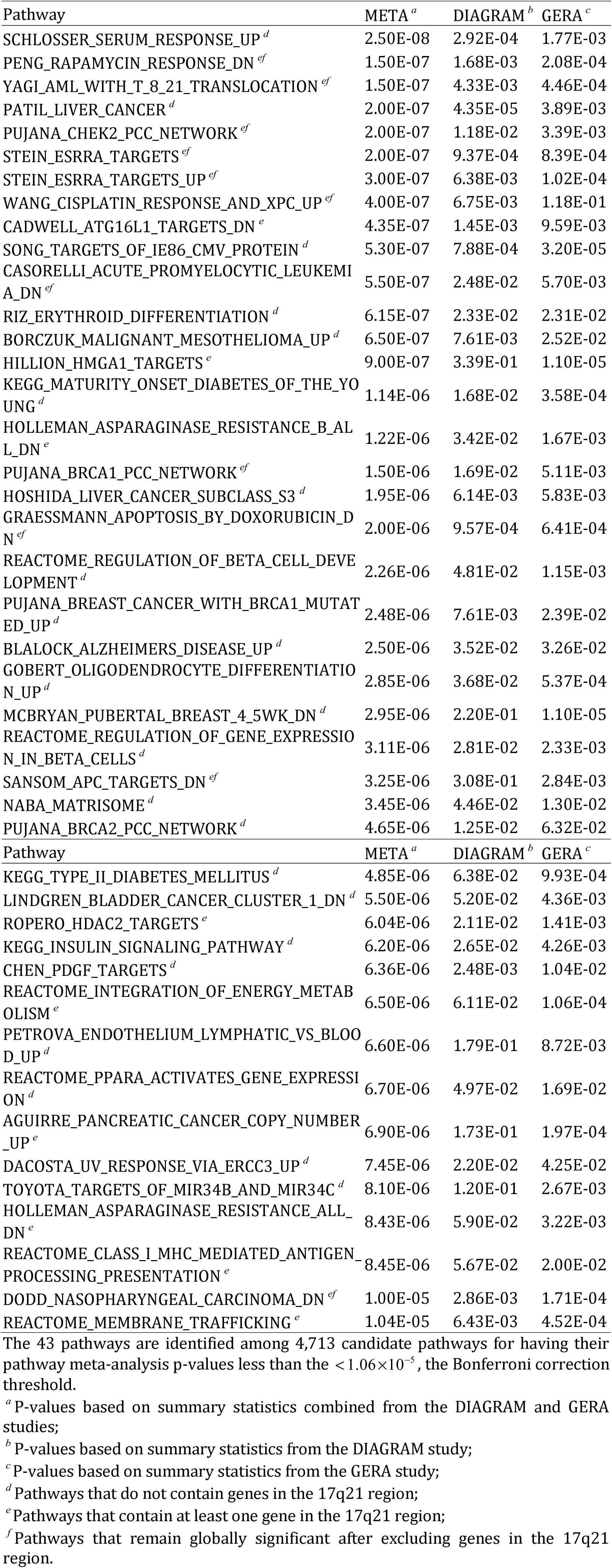
Summary of 43 significant pathways detected by the pathway meta-analysis based on the DIAGRAM and GERA studies

From Table 3, we can notice that some identified pathways have relatively weak association signals from each of the two studies, but have very significant p-values based on the pathway meta-analysis on the two studies combined. For example, the pathway RIZ_ERYTHROID_DIFFERENTIATION has p-values of 0.0233 and 0.0231 based on DIAGRAM and GERA studies, respectively. Combining these two p-values using Fisher’s method yields a p-value of 0.0046. On the other hand, the pathway meta-analysis produces a much more significant result (*p* = 6.15×10^−7^). This demonstrates the power advantage of the pathway meta-analysis over the approach that simply combines the pathways p-values from individual studies. The aforementioned simulation studies also confirmed this observation (Table 2).

In Figure 3, we illustrate the connection between the 43 significant pathways and a group of genes showing association evidence. For the purpose of illustration, in the figure we only focus on 46 genes that are covered by the 43 pathways and have their gene-level meta-analysis p-values less than 0.001. It is evident from Figure 3 that a cluster of 4 genes, *UBE2Z*, *SNF8*, *GIP*, and *ATP5G1*, has the most significant gene-level p-values (Table S3), and contribute association signals to 20 out of 43 significant pathways (Figures S6-S25). These 4 genes overlap each other at chromosome 17q21. This region contains a previously unidentified genome-wide significant synonymous SNP rsl058018 (meta-analysis *p* = 3.06 × 10^−8^) after two rounds of inflation adjustments. More detailed information on SNP rs1058018 and SNPs in that region are given in Table S4, Figure S4, and Figure S5. By conditioning on rsl058018, none of the other SNPs in this region are significant based on the conditional association analysis using the GERA individual-level GWAS data. Based on GTEx data v6, rsl058018 is a *cis* eQTL for *UBE2Z* in blood (*p* = 7.9×10^−15^). *UBE2Z* is involved in Class I MHC antigen processing and presentation (GeneCards). The region at 17q21 was previously implicated to be associated with T2D through a candidate gene/loci approach [36]. Although genes at the 17q21 region carry the strongest association signal, 11 out of those 20 pathways remain to be globally significant(*p* < 1.06×10^−5^) after excluding those genes from the pathway definition (Table 3).

**Figure 3.**
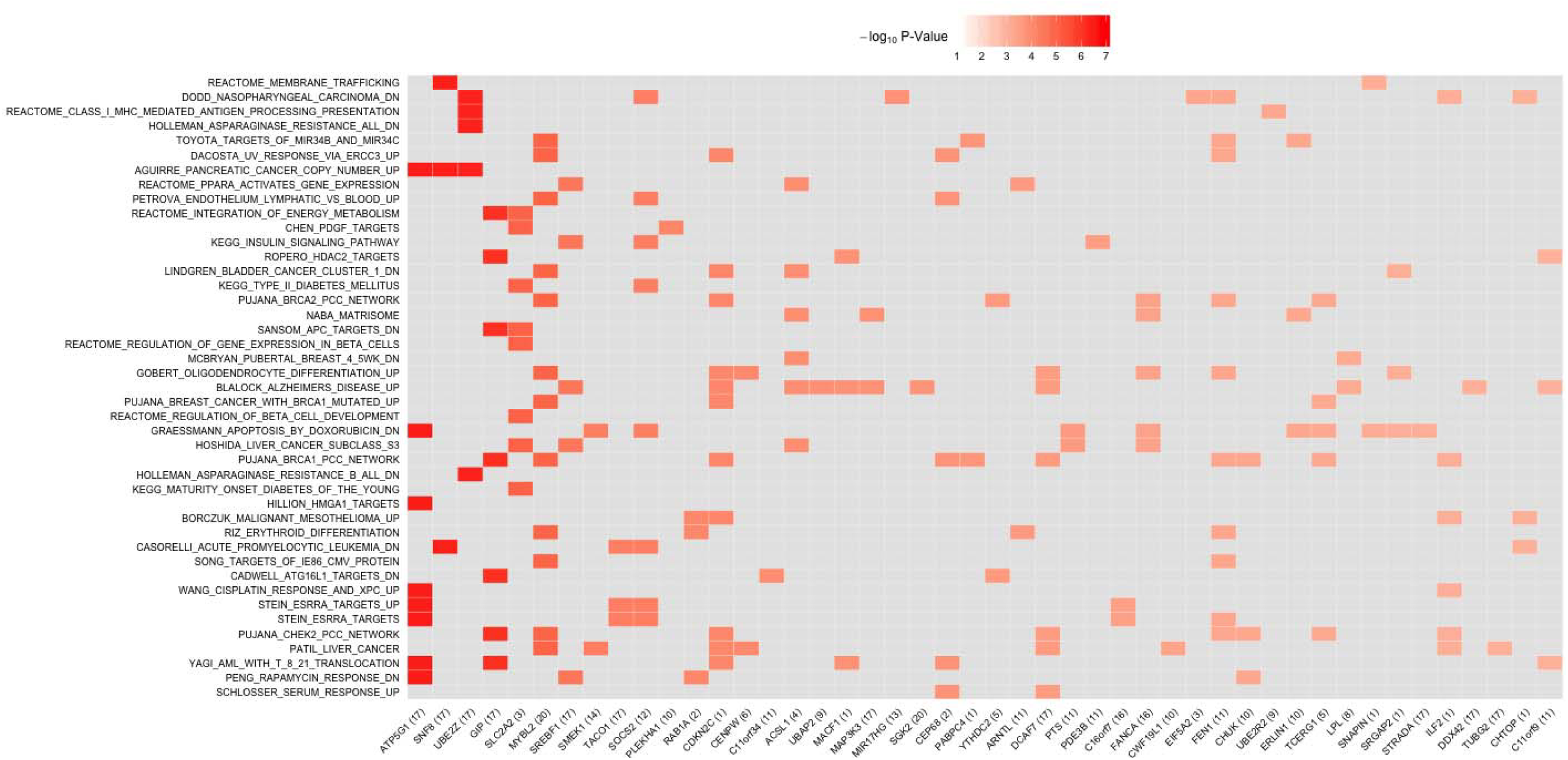
Heat map of gene-level p-values on selected genes within 43 significant pathways based on the DIAGRAM and GERA studies. There are 46 unique genes in the 43 significant pathways that have their gene-level meta-analysis p-values less than 0.001. Each row in the plot represents one of 43 significant pathways. Each column represents one of the 46 unique genes. The chromosome IDs of 46 unique genes are given in parentheses. The color of each cell represents the gene-level p-value (in the - log_10_ scale). A cell for a gene that is not included in a pathway is colored gray in the corresponding entry. The orders of genes (x-axis) and pathways (y-axis) are arranged according to their gene and pathway meta-analysis p-values.

The majority of 43 identified pathways are enriched with signals from multiple chromosomal regions as demonstrated by the Q-Q plots of their SNP-level and gene-level p-values (Figures S6-S48). For example, the strongest T2D-associated pathway, SCHLOSSER_SERUM_RESPONSE_UP, consists of 103 genes, which includes two genes with p-values < 0.001, and has 20 genes with p-values between 0.001 and 0.05 (Figure S26, and Supplemental data). We conducted the ingenuity pathway analysis on those 22 genes with p-values less than 0.05, and found enrichment of these genes in caveolae-mediated cytosis (important for removal of low/high density lipoproteins), and lipid metabolism pathways, and in functions/diseases related to differentiation of phagocytes and transport of proteins.

It is assuring that our pathway analysis detected several pathways that are natural candidates underlying the development of T2D, including the pathways KEGG_MATURITY_ONSET_DIABETES_OF_THE_YOUNG (Figure S31), KE GG_TYPE_II_DIABETES_MELLITUS (Figure S41), KEGG_INSULIN_SIGNALING_PATHWAY (Figure S43), and REACTOME_REGULATION_OF_BETA_CELL_DEVELOPMENT (Figure S33). It is worth emphasizing that these pathways were analyzed after excluding genes in the neighborhood of 170 GWAS established T2D loci and 195 SNPs with p-values < 10^−7^ on either DIAGRAM or GERA data, which suggests that these well-defined T2D-related pathways are enriched with additional unidentified and contributory T2D-associated genes.

Among the 43 globally significant pathways, there are multiple ones that are defined according to specific gene expression patterns on various tumor types, including pancreatic adenocarcinoma (Figure S21), hepatocellular carcinoma (HCC) (Figure S27, and S32), bladder carcinoma (Figure S42), nasopharyngeal carcinoma (Figure S24), and familial breast cancer (Figures S34, and S37). It is well recognized that T2D patients have elevated risk of cancer at multiple cancer sites, such as the liver and pancreas [37, 38]. These findings can provide valuable insights into the genetic basis underlying the connection between T2D and a host of different cancers.

In the above analysis, we used the sARTP method with the gene-level association evidence summarized by one or two most significant SNPs within each gene, under the assumption that there are at most two independent association signals within a given gene. We also applied sARTP by using 3 SNP-level cut points (i.e., (*c*_1_, *c*_2_, *c*_3_) = (1,2,3)) to reanalyze the 4,713 pathways based on the combined data of DIAGRAM and GERA. It appears that results obtained by sARTP with 3 SNP-level cut points are very consistent with those with 2 cut points (Figure S49).

#### Findings from Eastern Asian Populations

We reanalyzed the 43 significant pathways identified from the European populations using summary-level data generated by the AGEN-T2D study. An inflation factor *λ*_*GC*_ = 1.03 calculated from the AGEN-T2D meta-analysis was adjusted in the pathway meta-analysis. The genetic regions excluded from analyzing the DIAGRAM and GERA studies were also excluded from the AGEN study. The results were summarized in Table 4. There are 10 out of 43 pathways with the unadjusted p-value less than 0.05, suggesting that many pathways identified from the European populations were also enriched with T2D-associated genes in the eastern Asian populations. The Supplemental Data provides more details on results of those 10 pathways. Among the 43 pathways, we were able to identify 4 significant T2D-associated pathways at the false discovery rate (FDR [39]) of 0.05 (Figures S27, S12, S32 and S36), and 3 additional T2D-associated pathways at the FDR of 0.1 (Figures S47, S44, and S25). All the pathway p-values remain basically the same level if we further excluded genes within ±500kb regions surrounding the GWAS T2D loci established in eastern Asian populations. These results support the presence of trans-ethnic pathway effect on T2D in European and eastern Asian populations [11,12].

**Table 4.** Pathway p-values and FDR adjusted p-values based on the AGEN-T2D study.

| Pathway | P-value <sup>a</sup> | FDR <sup>b</sup> |
| --- | --- | --- |
| PATIL_LIVER_CANCER <sup>c</sup> | 0.0014 | 0.029 |
| CADWELL_ATG16L1_TARGETS_DN | 0.0023 | 0.029 |
| HOSHIDA_LIVER_CANCER_SUBCLASS_S3 <sup>c</sup> | 0.0025 | 0.029 |
| GOBERT_OLIGODENDROCYTE_DIFFERENTIATION_UP <sup>c</sup> | 0.0027 | 0.029 |
| DACOSTA_UV_RESPONSE_VIA_ERCC3_UP <sup>c</sup> | 0.011 | 0.074 |
| CHEN_PDGF_TARGETS <sup>c</sup> | 0.011 | 0.074 |
| REACTOME_MEMBRANE_TRAFFICKING | 0.012 | 0.074 |
| AGUIRRE_PANCREATIC_CANCER_COPY_NUMBER_UP | 0.026 | 0.14 |
| MCBRYAN_PUBERTAL_BREAST_4_5WK_DN <sup>c</sup> | 0.041 | 0.19 |
| LINDGREN_BLADDER_CANCER_CLUSTER_1_DN <sup>c</sup> | 0.043 | 0.19 |
| PUJANA_CHEK2_PCC_NETWORK | 0.057 | 0.21 |
| BLALOCK_ALZHEIMERS_DISEASE_UP <sup>c</sup> | 0.059 | 0.21 |
| SCHLOSSER_SERUM_RESPONSE_UP <sup>c</sup> | 0.085 | 0.27 |
| RIZ_ERYTHROID_DIFFERENTIATION <sup>c</sup> | 0.089 | 0.27 |
| DODD_NASOPHARYNGEAL_CARCINOMA_DN | 0.097 | 0.28 |
| TOYOTA_TARGETS_OF_MIR34B_AND_MIR34C <sup>c</sup> | 0.10 | 0.28 |
| REACTOME_CLASS_I_MHC_MEDIATED_ANTIGEN_PROCESSING_PRESENTATION | 0.14 | 0.32 |
| PUJANA_BRCA2_PCC_NETWORK<sup>c</sup> | 0.14 | 0.32 |
| WANG_CISPLATIN_RESPONSE_AND_XPC_UP | 0.14 | 0.32 |
| STEIN_ESRRA_TARGETS | 0.16 | 0.35 |
| PUJANA_BREAST_CANCER_WITH_BRCA1_MUTATED_UP<sup>c</sup> | 0.17 | 0.35 |
| GRAESSMANN_APOPTOSIS_BY_DOXORUBICIN_DN | 0.18 | 0.36 |
| PUJANA_BRCA1_PCC_NETWORK | 0.23 | 0.43 |
| HOLLEMAN_ASPARAGINASE_RESISTANCE_B_ALL_DN | 0.35 | 0.62 |
| PENG_RAPAMYCIN_RESPONSE_DN | 0.38 | 0.62 |
| ROPERO_HDAC2_TARGETS | 0.38 | 0.62 |
| KEGG_MATURITY_ONSET_DIABETES_OF_THE_YOUNG<sup>c</sup> | 0.39 | 0.62 |
| HILLION_HMGA1_TARGETS | 0.43 | 0.65 |
| HOLLEMAN_ASPARAGINASE_RESISTANCE_ALL_DN | 0.44 | 0.65 |
| PETROVA_ENDOTHELIUM_LYMPHATIC_VS_BLOOD_UP<sup>c</sup> | 0.45 | 0.65 |
| REACTOME_PPARA_ACTIVATES_GENE_EXPRESSION<sup>c</sup> | 0.52 | 0.72 |
| REACTOME_REGULATION_OF_GENE_EXPRESSION_IN_BETA_CELLS<sup>c</sup> | 0.55 | 0.74 |
| KEGG_INSULIN_SIGNALING_PATHWAY<sup>c</sup> | 0.58 | 0.74 |
| STEIN_ESRRA_TARGETS_UP | 0.59 | 0.74 |
| YAGI_AML_WITH_T_8_21_TRANSLOCATION | 0.64 | 0.76 |
| NABA_MATRISOME<sup>c</sup> | 0.65 | 0.76 |
| KEGG_TYPE_II_DIABETES_MELLITUS<sup>c</sup> | 0.67 | 0.76 |
| SANSOM_APC_TARGETS_DN | 0.69 | 0.76 |
| REACTOME_INTEGRATION_OF_ENERGY_METABOLISM | 0.70 | 0.76 |
| REACTOME_REGULATION_OF_BETA_CELL_DEVELOPMENT<sup>c</sup> | 0.70 | 0.76 |
| BORCZUK_MALIGNANT_MESOTHELIOMA_UP<sup>c</sup> | 0.75 | 0.79 |
| SONG_TARGETS_OF_IE86_CMV_PROTEIN<sup>c</sup> | 0.86 | 0.88 |
| CASORELLI_ACUTE_PROMYELOCYTIC_LEUKEMIA_DN | 0.88 | 0.88 |

1 These 43 pathways are nominated through the pathway meta-analysis on DIAGRAM 2 and GERA studies. The analysis is carried out on the summary data from the AGEN- 3 T2D study.
<sup>a</sup> P-values based on summary statistics from the AGEN-T2D study;
<sup>b</sup> FDR adjusted p-values;
<sup>c</sup> Pathways that do not contain genes in the 17q21 region.

Given the existing epidemiologic evidence on the close connection between T2D and the liver cancer, it is noteworthy that the two HCC related pathways (Figures S27,S32) identified in European populations remain to be significant in eastern Asian populations at the FDR of 0.05 (Table 4). The pathway PATIL_LIVER_CANCER consists of 653 genes (after data preprocessing) that are highly expressed in HCC and are enriched with genes having functions related to cell growth, cell cycle, metabolism, and cell proliferation [40]. The other pathway, HOSHIDA_LIVER_CANCER_SUBCLASS_S3 consists of 240 genes that show similar gene expression variation patterns and together define a HCC subtype with its unique histologic, molecular and clinical characteristics [41]. These two pathways have only 6 genes in common, and none of the 6 genes has a gene-level p-value < 0.05 in either European or eastern Asian data. More in depth investigations of these two complementary pathways could lead to further understanding the connection between T2D and the liver cancer.

The genome-wide significant SNP rs1058018 at the 17q21 region identified through the combined analysis of DIAGRAM and GERA studies turned out to be null in the AGEN-T2D study (*p* = 0.29). This could be due to the relatively small sample size of the AGEN-T2D study, or the genetic risk heterogeneity at the 17q21 locus among different ethnic populations. Nevertheless, 2 out of the 20 pathways (Figures S12 and S25) that contain genes within the 17q21 region are still significant at the FDR of 0.1. Among the 23 pathways that do not contain any gene within the 17q21 region, 5 pathways remain significant at the FDR of 0.1 (Figure S27, S32, S36, S47, and S44).

## Discussion

We developed a general statistical procedure sARTP for pathway analysis using SNP-level summary statistics generated from multiple GWAS. By applying sARTP to summary statistics from two large studies with a total of 19,809 T2D cases and 111,181 controls with European ancestry, we were able to identify 43 globally significant T2D-associated pathways after excluding genes in neighborhoods of GWAS established T2D loci. Using summary data generated from 8 T2D GWAS with 6,952 cases and 11,865 controls from eastern Asian populations, we further showed that 7 out of 43 pathways identified in the European populations were also significant in the eastern Asian populations at the FDR of 0.1. The analysis clearly highlights novel T2D-associated genes and pathways beyond what has been known from single-SNP association analysis reported from largest GWAS to date. Since the new procedure requires only SNP-level summary statistics, it provides a flexible way for conducting pathway analysis, alleviating the burden of handling large volumes of individual-level GWAS data.

We have developed a computationally efficient R package called ARTP2 implementing the ARTP and sARTP procedures, so that it can be used for conducting pathway analysis based on individual-level genetic data, as well as SNP-level summary data from one or multiple GWAS. The R package also supports the parallelization on Unix-like OS, which can substantially accelerate the computation of small p-values when a large number of resampling steps are needed. The ARTP2 package has a user-friendly interface and provides a comprehensive set of data preprocessing procedures to ensure that all the input information (e.g., allele information of SNP-level summary statistics and genotype reference panel) can be processed coherently. To make the sARTP method accessible to a wider research community, we have also developed a web-based tool that allows investigators to conduct their pathway analyses using the computing resource at the National Cancer Institutes through simple on-line inputs of summary data.

Single-locus analysis of GWAS usually has its genomic control inflation factor larger than 1.0. Some proportion of the inflation can be attributed to various confounding biases, such as the one caused by population stratification, while the other part can be due to the real polygenic effect. In the pathway analysis it is important to minimize the confounding bias at the SNP-level summary statistic. Otherwise a small bias at the SNP level can be accumulated in the pathway analysis, and lead to an elevated false discovery rate. Here we try to remove the confounding bias by adjusting for the genomic control inflation factor observed at the GWAS study. This approach is conservative because part of the inflation can be caused by the real polygenic effect. Recently, [42] developed the LD score regression method to quantify the level of inflation caused solely by the confounding bias. Adjusting for the inflation factor estimated by this method, instead of the genomic control inflation factor, can potentially increase the power of the pathway analysis. However, the LD score regression method relies on a specific polygenic risk model, and its estimate might not be robust for this model assumption. More investigations are needed to evaluate the impact of this new inflation adjustment on the pathway analysis.

There are several other strategies to increase the power of pathway analysis besides increasing sample size [4]. One area of active research is to find better ways to define the gene-level summary statistic using observed genotypes on multiple SNPs, so that it can accurately characterize the impact of the gene on the outcome [43–46]. In our proposed procedure, we adopt a data driven approach to select a subset of SNPs within a gene that collectively show the strongest association evidence. Because of this, we have to pay the penalty of multiple-comparison in the final pathway significance assessment. However, it is well recognized that SNPs at different loci can have varied levels of functional implications. We can potentially reduce the burden of multiple-comparisons and thus improve the power of the pathway analysis, by prioritizing SNPs according to existing genomic knowledge and other data resources. For example, [47] recently proposed a new gene-level summary statistic based on a prediction model that was trained with external transcriptome data. The gene-level summary statistic is defined as the predicted value that estimates the component of gene expression regulated by a subject’s genotypes within the neighborhood of the considered gene. Pathway analysis procedures using this kind of biologically informed gene-level summary statistic can be easily incorporated into the ARTP2 framework.

The sARTP method can be easily expanded to adopt other multi-locus statistics in accumulating association within a gene, as long as they can be written in terms of SNP-level score statistics and their variance-covariance matrix. For example, the current ARTP2 package provides the option for conducting the pathway meta-analysis using the joint test statistics proposed by [31].

When conducting pathway analysis with individual-level genetic data, we could run into a computing memory issue if the study has a large sample size and the pathway consists of a large number of genes and SNPs (Figure S4). The ability of performing pathway analysis using summary data provides a convenient and efficient solution in those situations. We can first calculate the SNP-level summary statistics based on the individual-level genetic data, and then randomly sample a small proportion of the original data as an internal reference to estimate the variance-covariant matrix for score statistics at considered SNPs. Based on our experiments, using 500 or more subjects to form a reference panel would be good enough to generate accurate pathway p-values. As shown in Figure 1, the testing results using this approach are very consistent with those based on individual-level genotype data.

The sARTP approach can be applied directly to SNP-level meta-analysis results. This is very convenient as meta-analysis results are in general easily accessible. But we want to emphasize that it is important to know the set of the SNPs studied by each participating study in order to apply sARTP properly, as the SNP coverage information is essential for accurately estimating the variance-covariance matrix of SNP-level score statistics. GWAS consortia usually do not post the SNP coverage information when releasing their meta-analysis results. Many statistical packages designed for conducting multi-locus analysis based on meta-analysis results often assume the uniform coverage [15-18, 24, 25, 48]. As we already have demonstrated in the context of pathway analysis, this type of over-simplification could lead to inflated false positive rate.

The proposed procedure assumes that all participating studies are conducted with subjects with the same ancestry background. If this is not the case, a simple approach is to use the Fisher’s method to combine pathway p-values estimated on different ethnic populations. However, if there were no evidence for the existence of cross ethnic risk heterogeneity, it would be more powerful to assume a fixed effects model on the SNP-level association when performing the pathway analysis. In that case, since the LD structures in different ethnic populations are different, we need a separate reference panel for each ethic group to derive the corresponding variance-covariance matrix of the score statistics. The current ARTP2 package needs to be modified to accommodate such a more complicated case.

As already demonstrated by many successful GWAS meta-analysis, increasing the sample size through combining results from multiple studies is a very effective way to improve our chance for new findings. For the same reason, pathway-based meta-analysis can provide us with new opportunities to uncover biological pathways that are previously undetectable due to the limitation on the sample size. With more summary data from meta-analysis becoming increasingly available, we expect the ARTP2 package would be a valuable tool for further exploring the genome in search for the hidden heritability.

## Appendix A: Recovering Score Statistics and Its Variance-Covariance Matrix Using Summary Results from the Fixed Effect Model

Here we derive the approximated score statistic *S* and its variance-covariance matrix *V* using summary statistics from the fixed effect model. Based on (4), it is straightforward to see that 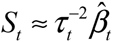. Note that *V*_*ts*_ in equation (5) depends on 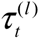 estimated from individual studies, which cannot be derived from *τ*_*t*_. However,assume that 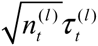 can be approximated as an unknown but common constant value *v*_*t*_ across all studies, and if *y*̅^(*l*)^(1 – *y*̅^(*l*)^) ≈ *y*̅(1 – *y*̅), we have 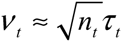 and 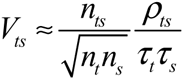. The similar argument has been used in Lin and Zeng (49) to demonstrate that the meta-analysis is as efficient as the pooled analysis under those conditions.

## Appendix B: Further Evaluation of sARTP Under the Null

We conducted additional experiments to evaluate the empirical size of sARTP. Based on the GERA T2D GWAS, we created 20 GWAS data under the null by randomly permuting the outcome, while keeping individual genotypes unchanged. On each null data, we excluded 274 pathways with over 10,000 SNPs for the sake of reducing computational burden, and conducted a pathway-based meta-analysis with sARTP on the remaining 4,439 pathways defined in MSigDB v5.0. The Q-Q plots of the pathway p-values of these 20 experiments are shown in Figure S50. Since there are extensive overlaps between pathways, their pathway p-values in each experiment are correlated. As a result, the Q-Q plot has a large variation around the diagonal line. But on average, there is no apparent genomic control inflation across 20 experiments. Based on those 20 experiments, Table S5 shows the genomic control inflation factors, Spearman’s rank correlation coefficient between the pathway size (in terms of the number of unique SNPs, or genes in a pathway) and its pathway p-value. By inspecting those correlation coefficients, we did not see any evidence suggesting that the association significance level of a pathway is influenced by its size under the null.

## Supporting Information

Table S1. Power comparison between sARTP and aSPUsPath under the scenrio where each outcome-associated gene contains one functional SNP

Table S2. Power comparison between sARTP and aSPUsPath under the scenrio where each outcome-associated gene contains one or two functional SNP(s) with equal probability

Table S3. Summary of top 50 genes with smallest gene-level p-values from the gene-level meta-analysis based on the DIAGRAM and GERA studies

Table S4. Effect of SNP rs1058018 on type 2 diabetes

Table S5. The genomic control inflation factors, Spearman’s rank correlation coefficients between the pathway size and its p-value based on results obtained by applying sARTP to 20 simulated GWAS under the null

Figure S1. The sample size used for the study of each SNP in the DIAGRAM meta-analysis

Figure S2: Histograms of numbers of SNPs and genes after SNP filtering within each of 4,718 pathways in pathway analyses of the DIAGRAM study, the GERA study, and the two studies combined (META)

Figure S3: Boxplot of the number of pathways containing genes with p-values in a given range

Figure S4. The LocusZoom plot showing ±100kb region of rsl058018 in European populations

Figure S5: The LocusZoom plot showing ±100kb region of rsl058018 in eastern Asian populations

Figure S6: Q-Q plots for SNP p-values and sARTP gene p-values of pathway PENG_RAPAMYCIN_RESPONSE_DN.

Figure S7: Q-Q plots for SNP p-values and sARTP gene p-values of pathway YAGI_AML_WITH_T_8_21_TRANSLOCATI0N

Figure S8: Q-Q plots for SNP p-values and sARTP gene p-values of pathway PUJANA_CHEK2_PCC_NETWORK

Figure S9: Q-Q plots for SNP p-values and sARTP gene p-values of pathway STEIN_ESRRA_TARGETS

Figure S10: Q-Q plots for SNP p-values and sARTP gene p-values of pathway STEIN_ESRRA_TARGETS_UP

Figure S11: Q-Q plots for SNP p-values and sARTP gene p-values of pathway WANG_CISPLATIN_RESPONSE_AND_XPC_UP

Figure S12: Q-Q plots for SNP p-values and sARTP gene p-values of pathway CADWELL_ATG16L1_TARGETS_DN

Figure S13: Q-Q plots for SNP p-values and sARTP gene p-values of pathway CASORELLI_ACUTE_PROMYELOCYTIC_LEUKEMIA_DN

Figure S14: Q-Q plots for SNP p-values and sARTP gene p-values of pathway HILLION_HMGA1_TARGETS

Figure S15: Q-Q plots for SNP p-values and sARTP gene p-values of pathway HOLLEMAN_ASPARAGINASE_RESISTANCE_B_ALL_DN

Figure S16: Q-Q plots for SNP p-values and sARTP gene p-values of pathway PUJANA_BRCA1_PCC_NETWORK

Figure S17: Q-Q plots for SNP p-values and sARTP gene p-values of pathway GRAESSMANN_APOPTOSIS_BY_DOXORUBICIN_DN

Figure S18: Q-Q plots for SNP p-values and sARTP gene p-values of pathway SANSOM_APC_TARGETS_DN

Figure S19: Q-Q plots for SNP p-values and sARTP gene p-values of pathway R0PERO_HDAC2_TARGETS

Figure S20: Q-Q plots for SNP p-values and sARTP gene p-values of pathway REACTOME_INTEGRATION_OF_ENERGY_METABOLISM

Figure S21: Q-Q plots for SNP p-values and sARTP gene p-values of pathway AGUIRRE_PANCREATIC_CANCER_COPY_NUMBER_UP

Figure S22: Q-Q plots for SNP p-values and sARTP gene p-values of pathway HOLLEMAN_ASPARAGINASE_RESISTANCE_ALL_DN

Figure S23: Q-Q plots for SNP p-values and sARTP gene p-values of pathway REACTOME_CLASS_I_MHC_MEDIATED_ANTIGEN_PROCESSING_PRESENTATION

Figure S24: Q-Q plots for SNP p-values and sARTP gene p-values of pathway DODD_NASOPHARYNGEAL_CARCINOMA_DN

Figure S25: Q-Q plots for SNP p-values and sARTP gene p-values of pathway REACTOME_MEMBRANE_TRAFFICKING

Figure S26: Q-Q plots for SNP p-values and sARTP gene p-values of pathway SCHLOSSER_SERUM_RESPONSE_UP

Figure S27: Q-Q plots for SNP p-values and sARTP gene p-values of pathway PATIL_LIVER_CANCER

Figure S28: Q-Q plots for SNP p-values and sARTP gene p-values of pathway SONG_TARGETS_OF_IE86_CMV_PROTEIN

Figure S29: Q-Q plots for SNP p-values and sARTP gene p-values of pathway RIZ_ERYTHROID_DIFFERENTIATION

Figure S30: Q-Q plots for SNP p-values and sARTP gene p-values of pathway BORCZUK_MALIGNANT_MESOTHELIOMA_UP

Figure S31: Q-Q plots for SNP p-values and sARTP gene p-values of pathway KEGG_MATURITY_ONSET_DIABETES_OF_THE_YOUNG

Figure S32: Q-Q plots for SNP p-values and sARTP gene p-values of pathway HOSHIDA_LIVER_CANCER_SUBCLASS_S3

Figure S33: Q-Q plots for SNP p-values and sARTP gene p-values of pathway REACTOME_REGULATION_OF_BETA_CELL_DEVELOPMENT

Figure S34: Q-Q plots for SNP p-values and sARTP gene p-values of pathway PUJANA_BREAST_CANCER_WITH_BRCA1_MUTATED_UP

Figure S35: Q-Q plots for SNP p-values and sARTP gene p-values of pathway BLALOCK_ALZHEIMERS_DISEASE_UP

Figure S36: Q-Q plots for SNP p-values and sARTP gene p-values of pathway GOBERT_OLIGODENDROCYTE_DIFFERENTIATION_UP

Figure S37: Q-Q plots for SNP p-values and sARTP gene p-values of pathway MCBRYAN_PUBERTAL_BREAST_4_5WK_DN

Figure S38: Q-Q plots for SNP p-values and sARTP gene p-values of pathway REACTOME_REGULATION_OF_GENE_EXPRESSION_IN_BETA_CELLS

Figure S39: Q-Q plots for SNP p-values and sARTP gene p-values of pathway NABA_MATRISOME

Figure S40: Q-Q plots for SNP p-values and sARTP gene p-values of pathway PUJANA_BRCA2_PCC_NETWORK

Figure S41: Q-Q plots for SNP p-values and sARTP gene p-values of pathway KEGG_TYPEJI_DIABETES_MELLITUS

Figure S42: Q-Q plots for SNP p-values and sARTP gene p-values of pathway LINDGREN_BLADDER_CANCER_CLUSTER_1_DN

Figure S43: Q-Q plots for SNP p-values and sARTP gene p-values of pathway KEGG_INSULIN_SIGNALING_PATHWAY

Figure S44: Q-Q plots for SNP p-values and sARTP gene p-values of pathway CHEN_PDGF_TARGETS

Figure S45: Q-Q plots for SNP p-values and sARTP gene p-values of pathway PETROVA_ENDOTHELIUM_LYMPHATIC_VS_BLOOD_UP

Figure S46: Q-Q plots for SNP p-values and sARTP gene p-values of pathway REACTOME_PPARA_ACTIVATES_GENE_EXPRESSION

Figure S47: Q-Q plots for SNP p-values and sARTP gene p-values of pathway DACOSTA_UV_RESPONSE_VIA_ERCC3_UP

Figure S48: Q-Q plots for SNP p-values and sARTP gene p-values of pathway TOYOTA_TARGETS_OF_MIR34B_AND_MIR34C

Figure S49. Comparison of sARTP p-values obtained with 2 or 3 SNP-level cut points based on combined data of DIAGRAM and GERA

Figure S50. Q-Q plots of pathway p-values based on 20 GWAS datasets generated under the null

## Acknowledgments

This study utilized the computational resources of the NIH HPC Biowulf cluster (https://hpc.nih.gov/). The authors thank the AGEN-T2D consortium and DIAGRAM consortium for sharing the meta-analysis summary data. The authors acknowledge Sue Pan, JJ Pan, Wesley Obenshain, Tony Hall, Jim Zhou, Ye Wu, Cuong Nguyen for their help in developing the web-based tool for ARTP2.

## Web Resources

The URLs for data and software presented herein are as follows:

DIAbetes Genetics Replication And Meta-analysis (DIAGRAMv3), http://diagramconsortium.org/

Genetic Epidemiology Research on Aging (GERA, dbGaP Study Accession: phs000674.v1.p1), http://www.ncbi.nlm.nih.gov/proiects/gap/cgibin/studv.cgi?study_id=phs000674.v1.p1

Molecular Signatures Database (C2: curated gene sets), http://software.broadinstitute.org/gsea/msigdb/collections.isp#C2

BioMart (Homo sapiens genes NCBI36 and GRCh37.p13), http://feb2014.archive.ensembl.org/

IMPUTE2, https://mathgen.stats.ox.ac.uk/impute/impute_v2.html

GWAS Catalog, http://www.ebi.ac.uk/gwas/

1000 Genomes Project (Phase 3, v5, 2013/05/02), ftp://ftp.1000genomes.ebi.ac.uk/vol1/ftp/release/20130502/

aSPU, https://cran.r-proiect.org/web/packages/aSPIJ/index.html

GTEx Portal v6, http://gtexportal.org/home/

GeneCards Human Gene Database, http://www.genecards.org/

Ingenuity Pathway Analysis, http://www.ingenuity.com/

LocusZoom, http://locuszoom.sph.umich.edu/locuszoom/

ARTP2 package, https://cran.r-project.org/web/packages/ARTP2/

Web-based tool of ARTP2, http://analysistools.nci.nih.gov/pathway/

